# Crucial role of Juvenile Hormone receptor components Methoprene-tolerant and Taiman in sexual maturation of adult male desert locusts (*Schistocerca gregaria*)

**DOI:** 10.1101/2020.09.01.277558

**Authors:** Michiel Holtof, Marijke Gijbels, Joachim van Lommel, Elfie Dekempeneer, Bart Nicolai, Jozef Vanden Broeck, Elisabeth Marchal

## Abstract

The desert locust, *Schistocerca gregaria*, is an infamous migrating insect, generally considered to be one of the most dangerous pest species worldwide. In an area that encompasses one fifth of the Earth’s land surface, desert locust swarms can devastate the agricultural production. Consequently, they pose a serious threat to food security and can have a dramatic socio-economic impact. Currently (in 2020), large parts of East Africa, Southwest and South Asia are experiencing the worst desert locust plague in many decades. Exceptionally high rainfall in different regions has caused the favorable environmental conditions for very successful reproduction and population growth. To better understand the molecular mechanisms responsible for this remarkable reproductive capacity, as well as to fill existing knowledge gaps regarding the regulation of male reproductive physiology, we decided to investigate the role of the *methoprene-tolerant (Scg-Met*) and *taiman (Scg-Tai*) genes in adult male desert locusts. Methoprene-tolerant and Taiman belong to the basic helix-loop-helix/Per-Arnt-Sim (bHLH/PAS) family of transcription factors and form a complex that is responsible for transducing the Juvenile Hormone (JH) signal. So far, *in vivo* research has mainly focused on the role of these JH receptor components in post-embryonic development, as well as in the reproductive physiology of adult female insects. In the current study, we show that silencing these components by RNA interference strongly inhibits sexual maturation of gregarious desert locust males, thereby severely disrupting reproduction. This observation is evidenced by the absence of a yellow colored cuticle, the significant reduction in the relative weight of the testes, the inability to show mating behavior, and the drastically reduced phenylacetonitrile (PAN) pheromone levels of the treated males. In addition, we also observed significant reductions of the relative weight, as well as the relative protein content, of the male accessory glands in *Scg-Met* knockdown locusts. Interestingly, in these animals the size of the *corpora allata* (CA), the endocrine glands where JH is synthesized, was significantly increased, and a significant rise was observed in the relative transcript levels of JH acid methyltransferase (JHAMT), a rate-limiting enzyme in the JH biosynthesis pathway. Moreover, other endocrine pathways also appeared to be affected by the knockdown, as evidenced by significant changes in the levels of the transcripts coding for the insulin-related peptide and for two neuroparsins in the fat body. Our results demonstrate that JH signaling pathway components play a crucial role in male reproductive physiology. Since they are crucial for a successful reproduction, they may hold great potential as candidate targets for the design of novel strategies for locust population management.

**Significance statement:** Desert locusts can form devastating swarms that threaten the livelihood in many of the world’s poorest countries, as demonstrated by the plagues in East Africa, Southwest and South Asia that are currently covered in the news worldwide. Control of this migratory pest mainly relies on the large-scale use of neurotoxic insecticides, also affecting non-target organisms in their already fragile habitats. Therefore, the identification of novel molecular targets for the design of more sustainable and more selective pest management techniques remains of crucial importance. Potential candidate targets in this respect are pathway components that transduce signals of typical insect hormones, such as ecdysteroids and juvenile hormones. By focusing in our current study on the JH receptor components, we showed that the reproductive capacities of desert locust males can be severely inhibited in an early stage, resulting in individuals that display no mating behavior, do not produce an important pheromone and do not show the characteristic yellow color of sexually mature gregarious males. Specifically targeting molecular components of the JH signaling pathway in future, more biorational control strategies could therefore be a very successful way by which expansion of these devastating plagues might be prevented.

## Introduction

Reproductive physiology in animals is strictly regulated by hormones. However, depending on the species, these hormones and their specific contributions may differ. This is illustrated nicely by reports on the endocrine regulation of the female reproductive physiology in a wide variety of insect species (as reviewed by Raikhel et al., 2005). Two important players in this process are the sesquiterpenoid, Juvenile Hormone (JH), and the ecdysteroid, 20-hydroxyecdysone (20E). Both lipophilic hormones are also key in the control of larval development and metamorphosis of insects. Whereas 20E triggers successive molts throughout the insect’s life cycle, JH, as its name indicates, represses metamorphosis and thus keeps the insect in a juvenile stage. Ecdysone is synthesized in the prothoracic glands of juvenile insects. These glands degenerate in adult insects and ecdysteroid synthesis takes place in the reproductive system (Brown et al., 2009; Marchal et al., 2010; Van de Velde et al., 2009). JH on the other hand is produced in the *corpora allata* (CA), small paired endocrine glands situated in the head (Marchal et al., 2010; Nation, 2002).

The 20E signaling pathway with its heterodimer of ecdysone receptor (EcR) and retinoid X receptor/ultraspiracle (RXR/USP) and downstream transcription factors, many belonging to the nuclear receptor superfamily, has been widely researched (Roy et al., 2018). However, for a long time the JH signaling pathway remained elusive. Methoprene-tolerant (Met), a member of the basic-helix-loop-helix (bHLH)/Per-Arnt-Sim (PAS) family of transcription factors, was first described in 1986 (Wilson and Fabian, 1986) as a crucial factor in the resistance to the commercially available JH analog, methoprene. The possibility that Met might act as a JH receptor was later confirmed by Miura et al. (Miura et al., 2005), who showed that the recombinant *Drosophila melanogaster* Met protein can bind JH with high affinity. The study by Charles et al. (Charles et al., 2011) later showed that a *Tribolium castaneum* Met contains a conserved hydrophobic binding pocket within the PAS-B domain, which binds JH and its analogs with high affinity, and that this capacity is necessary for interaction of Met with its partner, another member of the bHLH-PAS family named SRC (homologous to the mammalian steroid receptor coactivator 1) or Taiman (Tai), following FlyBase nomenclature. Tai had previously been shown in yeast two-hybrid studies to interact with Met (Li et al., 2011; Zhang et al., 2011). Further downstream of the Met-Tai complex, the JH signal was later found to be transduced by Krüppel-homolog 1 (Kr-h1), a C2H2 zinc-finger containing transcription factor (Minakuchi et al., 2008).

The identification of the JH receptor components has stimulated RNA interference (RNAi) mediated silencing studies in different insect species. These reverse genetics studies first showed that Met, its binding partner Tai, and their downstream transcription factor Kr-h1 are indeed transducing the anti-metamorphic JH signal (Coast and Schooley, 2011; Jindra et al., 2013; Konopova et al., 2011; Konopova and Jindra, 2007; Lozano et al., 2014; Parthasarathy et al., 2008). In both holo- and hemimetabolan insects it is now well-established that the anti-metamorphic action of JH is regulated by the MEKRE93 pathway, referring to the JH receptor Met and the transcription factors Kr-h1 and E93. Expression of E93, a key determinant promoting adult morphogenesis, is repressed by Kr-h1 during larval-to-larval transitions but will rise once JH titers drop and Kr-h1 expression is lowered before the onset of metamorphosis (Belles and Santos, 2014; Gujar and Palli, 2016; Ureña et al., 2014). Studies on Met, Tai and Kr-h1 were further elaborated towards other processes in which JH was reported to play a role, such as the developmental maturation of the female reproductive system and the reproductive diapause that is observed in a variety of insect species (Gujar and Palli, 2016; Liu et al., 2017; Marchal et al., 2014; Paim et al., 2012; Song et al., 2014; Villalobos-Sambucaro et al., 2015; J. L. Wang et al., 2017; Yue et al., 2018; Zhu et al., 2017). A recent study in the migratory locust, *Locusta migratoria*, has shown that one isoform of Tai contains a PRD-repeat motif that is essential for the induction of vitellogenesis by JH (Z. Wang et al., 2017). Until now, most research on the reproductive physiology of insects has focused on the role of these JH pathway components in females, while their exact role in males is much less understood.

Therefore, in the current study we decided to focus on the role of the JH receptor components Met and Tai in the reproductive maturation of males of the desert locust, *Schistocerca gregaria*. Maturation in crowd-reared adult male *S. gregaria* is associated with the display of sexual behavior and the bright yellow coloration of the cuticle due to the accumulation of carotenoids (Goodwin, 1952; Goodwin and Srisukh, 1948). The role of JH in male locusts has previously been studied by allatectomy (physical removal of the CA or chemical apoptosis of the CA cells using precocene) and JH (analog) treatments (Lange et al., 1983; Norris and Pener, 1965). In this regard, Amerasinghe (Amerasinghe, 1978) published an interesting study showing that JH can rescue the yellow coloration of the cuticle, as well as the production of maturation pheromone emissions in allatotectomized male *S. gregaria*. Phenylacetonitrile (PAN) later turned out to be a critical volatile in the pheromone bouquet of sexually mature gregarious male locusts (Amwayi et al., 2012; Loher, 1961; Mahamat et al., 2000). PAN acts as a courtship inhibiting pheromone with which sexually mature males can ensure postcopulatory mate guarding (Seidelmann and Ferenz, 2002). Although JH levels correlate well with PAN production in adult males, JH is suggested to regulate PAN biosynthesis only indirectly by stimulating the development of pheromone producing tissues (Ferenz and Seidelmann, 2003; Tawfik et al., 2000). Furthermore, a study by Braun and Wyatt (Braun and Wyatt, 1995) has shown that JH is necessary for the growth of the male accessory glands (AG) and for the associated protein synthesis in the fat body. The insect fat body plays a pivotal role in many metabolic processes, including nutrient and energy homeostasis, as well as reproduction (Arrese and Soulages, 2010). A clear correlation between the JH biosynthetic activity of the CA and the dry weight of the AG was also found in a study by Avruch and Tobe (Avruch and Tobe, 1978), who additionally showed a direct relationship between rates of JH release from the CA and the age of the animals. In the migratory grasshopper, *Melanoplus sanguinipes*, the biosynthesis of specific proteins in the AG is similarly highly dependent on JH (Ismail and Gillott, 1996).

Adequate nutrient uptake and allocation are pivotal for successful reproductive development in insects (Badisco et al., 2013). In general, the insulin/insulin-like growth factor (IGF) signaling pathway (ISP), sensing the organism’s nutrient status and positively controlling vitellogenesis and oocyte growth in female insects, plays a crucial role in the trade-off between reproduction and survival. Different studies have highlighted the complex interplay between JH and ISP in female reproductive physiology (Lenaerts et al., 2019; Mirth et al., 2019). In line with this, silencing of the insulin-related peptide (IRP) in female *S. gregaria* was shown to reduce vitellogenin (*Vg*) transcript levels and oocyte growth (Badisco et al., 2011). This study also showed that knocking down the *S. gregaria* neuroparsins (NPs), in contrast to *Scg-IRP*, increased *Scg-Vg* transcript levels and resulted in larger oocytes. NPs were initially characterized as antigonadotropic factors from the *pars intercerebralis-corpora cardiaca* (CC) neurohemal complex of locusts (Girardie et al., 1998, 1989, 1987). They belong to a large family of cysteine-rich secreted proteins, which also includes the ovary ecdysteroidogenic hormone (OEH), a gonadotropic factor in blood-fed mosquitoes, and shows sequence similarities with the most conserved region of vertebrate insulin-like growth factor binding proteins (IGFBPs) (Badisco et al., 2007, 2008; Nässel and Vanden Broeck, 2016). While in mosquitoes the gonadotropic OEH has been shown to act in parallel with the ISP via a Venus flytrap containing receptor tyrosine kinase, the exact mechanism(s) of *in vivo* action of the antigonadotropic *Scg-NPs* remain unknown (Lenaerts et al., 2017; Vogel et al., 2015).

In this paper, we report on the systemic RNAi knockdown of the JH receptor components in adult *S. gregaria* males. This experimental treatment significantly inhibited sexual maturation of male locusts, indicating that the molecular components of the JH receptor complex are crucial for this process. Our study shines new light on the molecular patterns of male desert locust maturation, a promising, but as yet largely neglected field of study in the search towards novel strategies to control devastating locust plagues.

## Material and methods

### Animal rearing

Desert locusts were reared under crowded conditions in large cages, in which temperature (32°C), relative humidity (40-60%) and light exposure (13 h photophase) were controlled. The animals were fed daily with dry oat flakes and fresh cabbage. Following mating, mature females were allowed to deposit their eggs in pots filled with damp sand. Each week, these pots were collected and transferred to empty cages, where eggs were allowed to hatch. In the described experiments, fifth nymphal and adult locusts were collected at the time of ecdysis and transferred to separate cages to obtain cohorts of temporally synchronized animals.

### *In silico* detection of JH signaling components

A BLAST search in an in-house whole-body transcriptome database of *S. gregaria* using publicly available (NCBI) sequences of Methoprene-tolerant and Taiman was performed to identify putative *S. gregaria* Met and Tai isoforms. Candidate nucleotide sequences were translated to the best candidate coding sequences (CDS) by Transdecoder. Coding regions were (1) scanned (HMMER) in the Pfam database, (2) scanned for potential signal peptide sequences with SignalP, (3) scanned for potential transmembrane domain sequences with TMHMM.

### Molecular cloning and sequencing of *Scg-Met* and *Scg-Tai*

Full length sequences for *Scg-Met* and *Scg-Tai* were found in an in-house whole-body transcriptome database of *S. gregaria*. The complete ORFs were PCR-amplified using *S. gregaria* cDNA originating from head, fat body and gonad tissues and Q5^®^ High-Fidelity DNA Polymerase (New England Biolabs, Ipswich, Massachusetts). Primers used for cloning are listed in Suppl. Table S2. The PCR products were run on a 1.2% agarose gel containing GelRed™ (Biotium, Fremont, California) to confirm their length. After electrophoresis, only a single band was observed, which was cloned (TOPO^®^ TA cloning kit for sequencing, Invitrogen) and sequenced (Sanger sequencing, LGC Genomics, Berlin, Germany) for verification.

### RNA interference

#### Production of dsRNA

dsRNA constructs for *Scg-Met* (Genbank: MK855050) (*dsScg-Met*) and *Scg-Tai* (Genbank: MK442071) (*dsScg-Tai*) were prepared using the MEGAscript^®^ RNAi Kit (ThermoFisher Scientific, Waltham, Massachusetts), designed for the production of dsRNAs longer than 200 bp, according to the manufacturer’s instructions. The procedure is based on the high-yield *in vitro* transcription reaction catalyzed by T7 RNA Polymerase starting from a user-provided linear template DNA flanked by T7 promoter sites. The dsRNA is obtained by annealing the resulting sense and antisense RNA strands. Primer sequences used for obtaining the linear templates are given in Suppl. Table S1. A PCR with REDTaq^®^ DNA polymerase (Sigma-Aldrich, Saint Louis, Missouri) was performed and the length of the amplified products was checked by 1% agarose gel electrophoresis. These bands were further cloned and sequenced (TOPO^®^ TA cloning kit for sequencing, Invitrogen) to confirm the amplicon sequence. For the production of GFP dsRNA (*dsGFP*), a PCR fragment flanked by a T7 promoter site was cloned in both sense and antisense direction in a TOPO 4.1 sequencing vector (Life Technologies, Carlsbad, California) and subsequently used as template for *in vitro* transcription. The final concentration of the produced dsRNA was estimated using a NanoDrop ND-1000 UV-VIS Spectrophotometer, and 1% agarose gel electrophoresis was performed to assess the integrity of the dsRNA.

#### RNAi experiment

To investigate the role of *Scg-Met* and *Scg-Tai* in the male reproductive physiology, adult male locusts were injected with 1 µg dsRNA (in 6 µL *S. gregaria* Ringer solution) targeting *Scg-Met* or *Scg-Tai* (1 L Ringer solution: 8.766 g NaCl; 0.188 g CaCl_2_; 0.746 g KCl; 0.407 g MgCl_2_; 0.336 g NaHCO_3_; 30.807 g sucrose; 1.892 g trehalose; pH 7.2). A control group was simultaneously injected with 1 µg *dsGFP*. Injections were initiated at the first day after the adult molt (day 0) and were repeated every 3 days to ensure a potent and persistent knockdown of the targets. The injections were pursued until day 34 after the adult molt. Then animals were sacrificed to collect their testes, AG, CA and fat body.

### Calculation of the accessory gland somatic and gonadosomatic indices

On day 34 of the adult stage, the live animals, as well as their dissected accessory glands (AG) and testes, were weighed using a Sartorius WME5004-e04112 balance. The accessory gland somatic (AGSI) and gonadosomatic (GSI) indices were calculated as the ratio of the AG and testes weight to the whole body weight, respectively. The protein content of the AG was determined using the bicinchoninic acid assay (BCA) method and normalized to the respective AG weight.

#### *Corpora allata* surface area measurement

The CA were carefully dissected and cleaned in *S. gregaria* Ringer solution and transferred to a small petri dish. Images of the CA were obtained using a light microscope (Zeiss SteREO Discovery.V8) equipped with an AxioCam ICc3 camera using the AxioVision 4.7 (Carl Zeiss, Oberkochen, Germany). These images were further processed in ImageJ allowing the surface area measurement of the CA in the image (Rasband, 2014) which was normalized to the locust’s total body weight.

### Volatile determination

On day 26 of the adult stage, per treatment 5 live locusts were analyzed for aroma. Individual locusts were introduced into 60 mL screw neck headspace vials (Macherey-Nagel, Dueren, Germany). The vial was sealed with TPFE 3.2 mm Beige screw cap (Marcherey-Nagel). The vial was incubated at 40 °C for 60 min. Then, volatiles were extracted with a DVB-CAR-PDMS fiber (Sigma-Aldrich) at 40 °C for 60 min. Determination of volatile compounds was performed on an Agilent 7890 A gas chromatograph (GC) (Agilent Technologies, USA) coupled to an Agilent 5975C VL MSD Mass Selective Detector (MS) (Agilent Technologies, Santa Clara, California) and equipped with a Gerstel Multipurpose Sampler 2 (Gerstel GmbH & Co.KG, Mülheim an der Ruhr, Germany). After extraction, aroma compounds were thermally desorbed into the injector heated at 220 °C and equipped with a SPME liner (0.75 i.d., Sigma-Aldrich). Split injection was performed in splitmode of 1:100 and the fiber thermally conditioned for 5 min. Separation was done on an HP-5MS column (30 m x 0.25 mm i.d. x 0.25 μm df) using helium as the carrier gas with a constant flow rate of 1 mL min-1. The column oven temperature program was as follows: 40 °C (4 min), 240 °C (10 °C min-1) and hold 240 °C (10 min). The total GC run time was 34 min. Mass spectra in the 35 to 350 m/z range were recorded at a scanning speed of 4.17 scan cycles per second. Chromatograms and mass spectra obtained from the GC-MS were deconvoluted and analyzed using MSD Chemstation (Agilent Technologies) and the automated mass spectral deconvolution and identification system (AMDIS) software v.2.1 (National Institute of Standards and Technology (NIST), Gaithersburg, Maryland). Aroma volatile compounds were identified by matching with NIST 11 mass spectral library. Volatile composition was compared by using absolute peak areas.

### Transcript *profiling*

#### Sample collection

Tissues (testes, AG, CA and fat body) were dissected in *S. gregaria* Ringer solution under a binocular microscope, and immediately transferred to liquid nitrogen to prevent RNA degradation. For each tissue and condition, five pooled samples were prepared as biological replicates. The tissue material collected in every pooled sample was derived from four individual locusts. These were stored at −80°C until further processing prior to RNA extraction.

#### RNA preparation

The pooled samples were transferred to MagNA Lyser Green Beads containing tubes and homogenized using a MagNA Lyser instrument (Roche, Basel, Switzerland). Total RNA was subsequently extracted from the tissue homogenate with the RNeasy Lipid Tissue Kit (Qiagen, Hilden, Germany) with additional DNase treatment according to the manufacturer’s instructions. Because of their relatively small size, CA were extracted using the RNAqueous-Micro Kit (Life Technologies), followed by the recommended DNase step. Quality and concentration of the resulting RNA samples were analyzed using a Nanodrop spectrophotometer (ThermoFisher Scientific).

#### Quantitative real-time (q-)RT-PCR

An equal amount of RNA was reverse transcribed into cDNA using the PrimeScript™ RT Reagent Kit, by following the manufacturer’s protocol (Takara, Shiga, Japan). Prior to q-RT-PCR transcript profiling, several previously described housekeeping genes (Van Hiel et al., 2009) were tested for their stability in the designed experiment. Optimal reference genes were selected using geNorm software (Vandesompele et al., 2002). q-RT-PCR primers for reference genes and target genes were designed using Primer Express software (Applied Biosystems, Foster City, California). For all investigated tissues and conditions, *Scg-Act* and *Scg-Ef1a* were selected by the geNorm software as having the most stable expression and were used as reference genes throughout this study. Primer sets were validated by designing relative standard curves with a serial ten-fold dilution of a calibrator cDNA sample. Efficiency of q-RT-PCR and correlation coefficient (R^2^) were measured for each primer pair. Primers for q-RT-PCR are given in Suppl. Table S2. All PCR reactions were performed in duplicate in 96-well plates on a StepOne System (ABI Prism, Applied Biosystems). Each reaction contained 5 μL fast Sybr Green, 0.5 μL of Forward and Reverse primers (10 μM), and 4 μL of cDNA. For all q-RT-PCR reactions, the following thermal cycling profile was used: 50 °C for 2 min, 95 °C for 10 min, followed by 40 cycles of 95 °C for 15 s and 60 °C for 60 s. Finally, a melting curve analysis was performed to check for primer dimers. For all transcripts, only a single melting peak was found during the dissociation protocol. Additionally, PCR products were run on a 1.2% agarose gel containing GelRed™ (Biotium). After electrophoresis only a single band was observed, which was cloned (TOPO^®^ TA cloning kit for sequencing, Invitrogen) and sequenced (Sanger sequencing, LGC Genomics, Berlin) to confirm target specificity. All q-RT-PCR results were normalized to the transcript levels of the selected reference genes and calculated relative to the transcript level in a calibrator sample according to the comparative Ct method (Vandesompele et al., 2002). GraphPad Prism 5 (GraphPad Software, San Diego, California) was used to test the statistical significance of differences in gene expression levels.

### q-RT-PCR temporal profiling of JH pathway components

For the temporal profiling of *Scg-Met, Scg-Tai, Scg-Krh1, Scg-JHAMT* and *Scg-CYP15*, q-RT-PCR analyses were performed on an in-house library of samples derived from adult male *S. gregaria* tissues. The investigated tissues in this library were the fat body, the gonads, the accessory glands and the CA/CC collected from male locusts at different timepoints during their adult development. This library was stored at −80°C and consisted of four pooled samples, each derived from five animals, for every tissue and time point. For these q-RT-PCR analyses, *Sg-Act* and *Scg-Ef1a* were selected by the geNorm software as having the most stable expression and were used as reference genes.

## Results

### Identification of *Scg-Met* and *Scg-Tai* cDNAs in *S. gregaria*

We identified the cDNAs coding for one Met isoform (*Scg-Met*) (Genbank: MK855050) and one Tai isoform (*Scg-Tai*) (Genbank: MK442071) by a BLAST search on an in-house whole-body transcriptome database of *S. gregaria* (unpublished data). Both sequences were uploaded to the NCBI GenkBank database. Both predicted proteins contain all necessary functional domains. The *Scg*-Tai sequence contains the basic helix-loop-helix (bHLH) and Per-Arnt-Sim (PAS) protein motifs typical of Tai proteins. The *Scg*-Met sequence contains the bHLH, PAS A, PAS B and PAC protein motifs inherent to Met proteins. Both sequences were validated. The identification of *Scg-Met* and *Scg-Tai* cDNAs allowed us to evaluate the role of these JH signal transducing factors in the reproductive development of adult *S. gregaria* males by means of RNAi-mediated knockdown.

### Transcript levels of JH pathway components in untreated adult male locusts

Until today, documentation on the role of JH signaling pathway components in the reproductive development of male insects is scarce. Therefore, we first analyzed the expression profiles of key mediators of this pathway in several relevant tissues at physiologically relevant time points during adult development of male desert locusts (Suppl. Fig. S1). The transcript levels of *Scg-Met, Scg-Tai, Scg-Krh1*, the transcription factor acting downstream of the JH receptor complex, and of juvenile hormone acid O-methyltransferase (*Scg*-JHAMT) and methyl farnesoate epoxidase (*Scg*-CYP15A1), the enzymes catalyzing the final steps in the JH biosynthetic pathway of the desert locust (Marchal et al., 2011), were analyzed in the fat body, the testes, the AG and the retrocerebral *corpora allata/corpora cardiaca* (CA/CC) complex of male adult locusts on day 3, day 7 and day 15 after their final molt. Within this timeframe, the male reproductive system fully develops, and the adult desert locusts gradually become sexually mature.

Transcript levels of *Scg-Met, Scg-Tai* and *Scg-Krh1* significantly increased from day 3, over day 7, to day 15 in the fat body of male desert locusts in the adult life stage (Suppl. Fig. S1). However, no significant changes in the levels of these transcripts were detected in either the testes or the AG.

In the CA/CC complex of male desert locusts, the relative transcript levels of *Scg-JHAMT* and *Scg-CYP15A1* significantly increased from day 3 to days 7 and 15 of the adult stage (Suppl. Fig. S1). Remarkably, in these CA/CC complexes the transcript levels of *Scg-Met* and *Scg-Krh1* were significantly lower on days 7 and 15, when compared to day 3.

### Effective silencing of the JH signaling pathway upon *dsScg-Met* injection

The efficiency of the RNAi-mediated knockdown of *Scg-Met* was investigated by comparing the transcript levels of *Scg-Met* in the fat body of *dsScg-Met* and *dsGFP* treated locusts on the day of dissections, 34 days after the adult molt. Our data clearly indicate the significant reduction of *Scg-Met* transcript levels, by an average of 44%, in the fat body of *dsScg-Met* injected locusts, confirming the effectiveness of the induced knockdown (p = 0.048) (Fig. 1 A). Since day 12 after the adult molt, all *dsGFP* injected (control) males demonstrated normal copulation behavior. However, during the entire timeframe of the experiment (34 days), none of the *dsScg-Met* injected (experimentally treated) animals had displayed any sexual activity. Thirty-four days after the adult molt, we investigated alterations in the levels of several transcripts coding for key proteins of the JH signaling pathway as a consequence of the *dsScg-Met* injections by means of q-RT-PCR. The mRNA levels of the downstream factor *Scg*-Krh1, generally considered as an important indicator of the Met-mediated JH signaling activity, were significantly reduced in both the fat body (p = 0.0021) and the CA (p = 0.0079) of *dsScg-Met* injected adult males, when compared to the *dsGFP* injected control animals (Fig. 1 B and Fig. 2 A). On the other hand, we detected a significant increase of the mRNA levels of the transcription factor *Scg*-E93 (p = 0.0079) and of the JH biosynthetic enzyme *Scg*-JHAMT (p = 0.008) in the CA of *dsScg-Met* injected males compared to control (*dsGFP*) locusts, while an upward (but non-significant) trend was observed for the *Scg-CYP15A1* transcript levels (p = 0.06) (Fig. 2 A).

**Figure 1.**
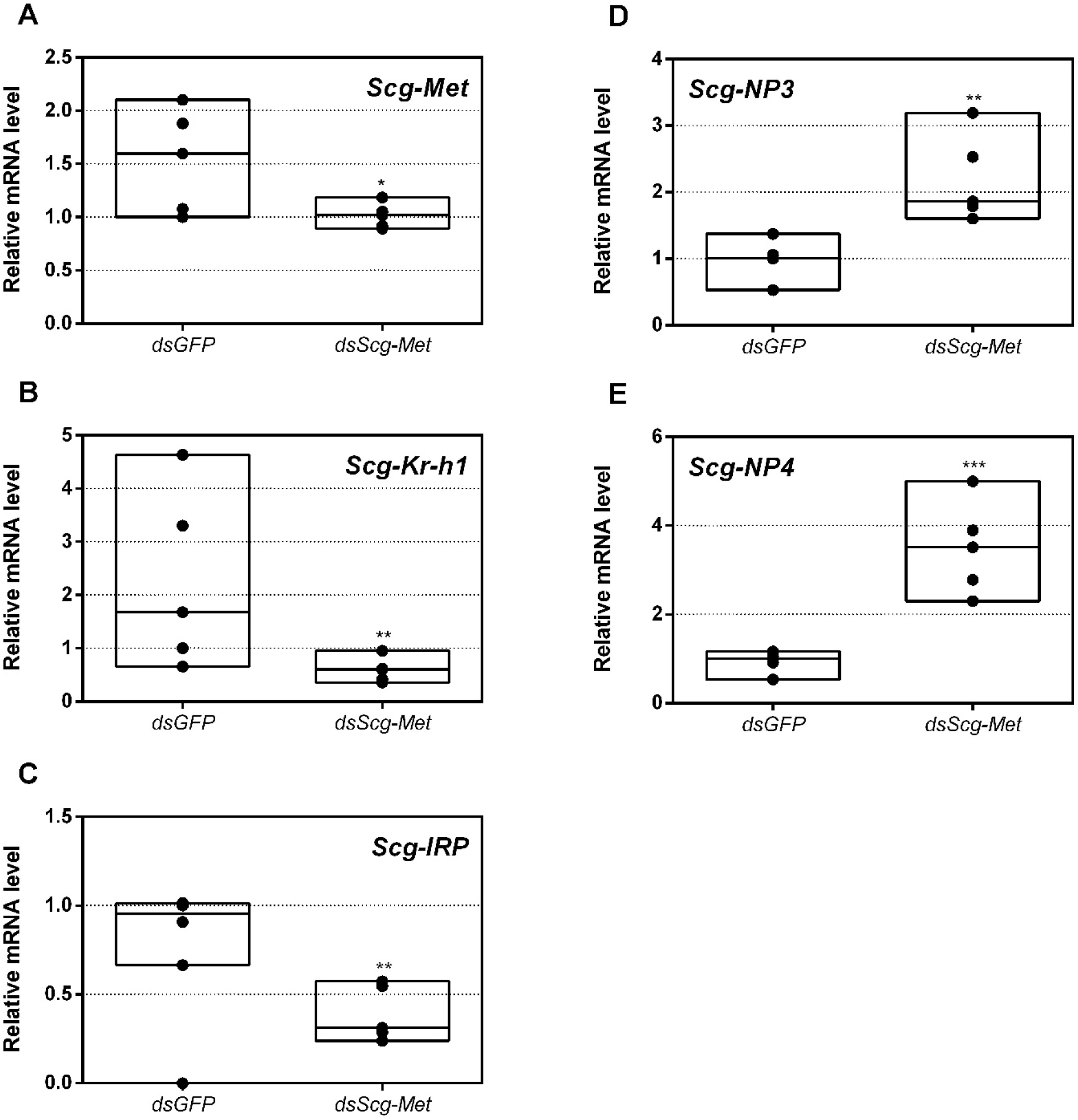
Relative mRNA levels measured in fat body tissue of *dsGFP* and *dsScg-Met* injected adult male desert locusts 34 days after the adult molt. **(A)** Transcript levels of *Scg-Met* were significantly reduced in the fat body demonstrating that the RNAi mediated knockdown was successful. Transcript levels of **(B)** *Scg-Krh1* and **(C)** *Scg-IRP* were significantly reduced, while transcript levels of **(D)** Scg-NP3 and **(E)** *Scg-NP4* were significantly increased in the fat body upon *dsScg-Met* treatment. Transcript levels in the fat body were normalized against two reference genes, *Scg-Act* and *Scg-Ef1a*. For every condition five biological replicates, each consisting of the pooled fat body samples from four individual locusts, were analyzed by q-RT-PCR. Data points are represented in a column graph as a floating bar (min to max value) with a line indicating the median. Significant differences (Student’s T-test) are indicated with asterisks (* p < 0.05; ** p < 0.01; *** p < 0.001). The following statistical p-values were obtained: *Scg-Met* (p = 0.048), *Scg-Krh1* (p = 0.0021), *Scg-IRP* (p = 0.0021), *Scg-NP3* (p = 0.0059), *Scg-NP4* (p = 0.0007). **Abbreviations:** ds = double stranded, *Scg = Schistocerca gregaria*, GFP = Green Fluorescent Protein, Met = Methoprene-tolerant, Kr-h1 = Krüppel-homolog 1, IRP = insulin-related peptide, NP = neuroparsin, Act = actin, Ef1a = elongation factor 1-alpha.

**Figure 2.**
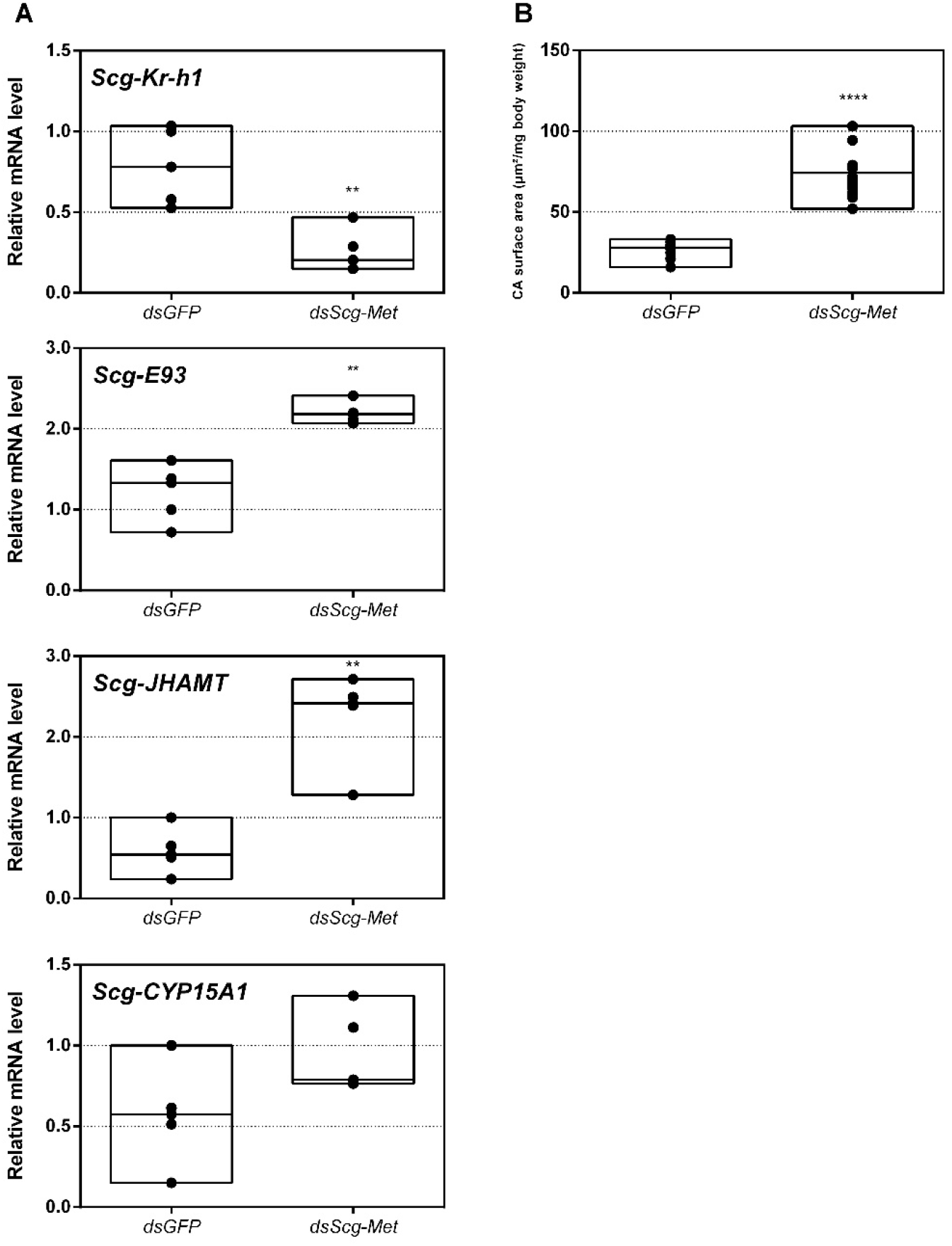
**(A)** Relative mRNA levels of *Scg-Krh1, Scg-E93, Scg-JHAMT* and *Scg-CYP15A* measured in the *corpora allata* (CA) of *dsGFP* and *dsScg-Met* injected adult male desert locusts 34 days after the adult molt. Data highlight the reduced *Scg-Krh1* transcript levels, as well as the increased *Scg-E93* and *Scg-JHAMT* transcript levels, in the CA after *Scg-Met* silencing. No significant change in the transcript levels of *Scg-CYP15A1* was detected. Transcript levels in the CA were normalized against two reference genes, *Scg-Act* and *Scg-Ef1a*. Data points are represented in a column graph as a floating bar (min to max value) with a line indicating the median. For every condition five biological replicates, each consisting of the pooled CA samples from four individual locusts, were analyzed by q-RT-PCR. Significant differences (Mann-Whitney U-test) are indicated with asterisks (** p < 0.01; **** p < 0.0001). The following statistical p-values were obtained: *Scg-Kr-h1* (p = 0.0079), *Scg-E93* (p = 0.0079), *Scg-JHAMT* (p = 0.008), *Scg-CYP15A1* (p = 0.06). **(B)** Normalized surface area of the CA in µm^2^ /mg body weight of *dsGFP* and *dsScg-Met* treated animals. These data demonstrate that the CA significantly increased in size after *dsScg-Met* treatment. On average the normalized surface area of the CA in control animals was 26.83 µm^2^/mg body weight, while the surface of the CA in *dsScg-Met* treated animals was 73.98 µm^2^/mg body weight. Data points are represented in a column graph as a floating bar (min to max value) with a line indicating the median. For each condition 13 animals were dissected for their CA. Significant differences (Student’s T-test) are indicated with asterisks (**** p < 0.0001). **Abbreviations:** ds = double stranded, *Scg = Schistocerca gregaria*, GFP = Green Fluorescent Protein, Met = Methoprene-tolerant, Kr-h1 = Krüppel-homolog 1, JHAMT = JH acid methyltransferase, CA = *corpora allata*, Act = actin, Ef1a = elongation factor 1-alpha.

### *Scg-Met* knockdown increases *corpora allata* size and JH biosynthetic enzyme expression

Upon dissection, 34 days after the adult molt, we observed that the CA of *dsScg-Met* injected animals were larger than the CA of control animals (p < 0.0001) (Fig. 2 B). On average the normalized surface area of the CA in control animals was 26.83 µm^2^/mg body weight, while the surface of the normalized surface are of the CA in *dsScg-Met* treated animals was 73.98 µm^2^/mg body weight. In addition, we also detected significantly increased *Scg-E93* (p = 0.0079) and *Scg-JHAMT* (p = 0.008) transcript levels in the CA upon Met knockdown (Fig. 2A).

### *Scg-Met* knockdown reduces yellow pigmentation in the cuticle of adult male desert locusts

While all control animals turned bright yellow, silencing of *Scg*-Met resulted in the absence of this yellow pigmentation (Fig. 3 B). Upon dissection, 34 days after the adult molt, the relative quantity of *Scg-YP* transcript, which codes for the carotenoid-binding Yellow Protein (Sas et al., 2007), was significantly lower in the epidermis of *dsScg-Met* injected males, when compared to the *dsGFP* injected ones (p = 0.0002) (Fig. 3A).

**Figure 3.**
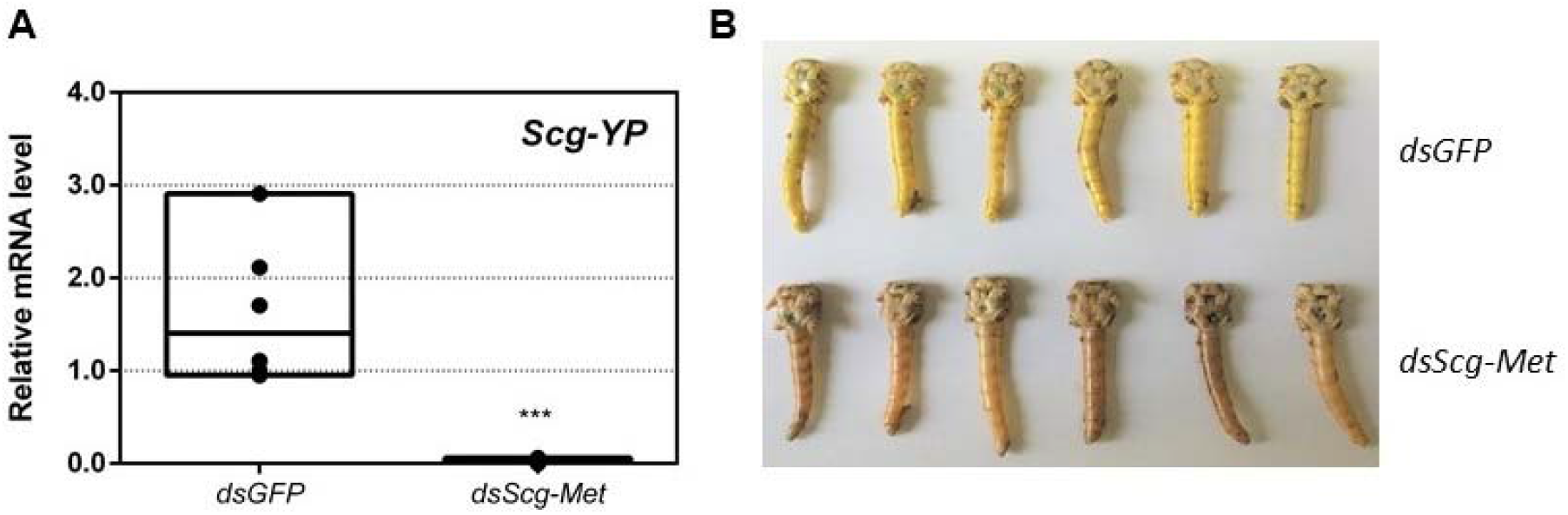
**(A)** Relative mRNA levels of *Scg-YP* measured in the epidermis of *dsGFP* and *dsScg-Met* injected adult male desert locusts 34 days after the adult molt. *Scg-YP* transcript levels were significantly lower after Met silencing. Transcript levels in the epidermis were normalized against two reference genes, *Scg-Act* and *Scg-Ef1a*. Data points are represented in a column graph as a floating bar (min to max value) with a line indicating the median. For every condition five biological replicates, each consisting of the pooled epidermis samples from four individual locusts, were analyzed by q-RT-PCR. Significant differences (Student’s T-test) are indicated with asterisks (*** p < 0.001) (the statistical p-value obtained for Scg-YP was 0.0002). **(B)** Pictures of the abdomen of both *dsGFP* treated (top, yellow) and *dsScg-Met* treated (bottom, beige/brown) male locusts 34 days after the adult molt. In clear contrast with the *dsScg-Met* injected locusts, all control animals displayed normal copulation behavior and had the bright yellow cuticular coloration that is typical for sexually mature gregarious desert locust males. **Abbreviations:** ds = double stranded, *Scg = Schistocerca gregaria*, GFP = Green Fluorescent Protein, Met = Methoprene-tolerant, YP = yellow protein, Act = actin, Ef1a = elongation factor 1-alpha.

### Effect of *Scg-Met* knockdown on phenylacetonitrile production

On day 26 of the adult stage, gas chromatography–mass spectrometry (GC–MS) was used to compare the PAN pheromone emission between *dsGFP* (control group) and *dsScg-Met* injected (experimental group) animals. These measurements demonstrated a very drastic reduction of PAN emission when *Scg-Met* was silenced (p < 0.0001)(Fig. 4). In contrast, the control animals showed normal PAN emission spectra.

**Figure 4.**
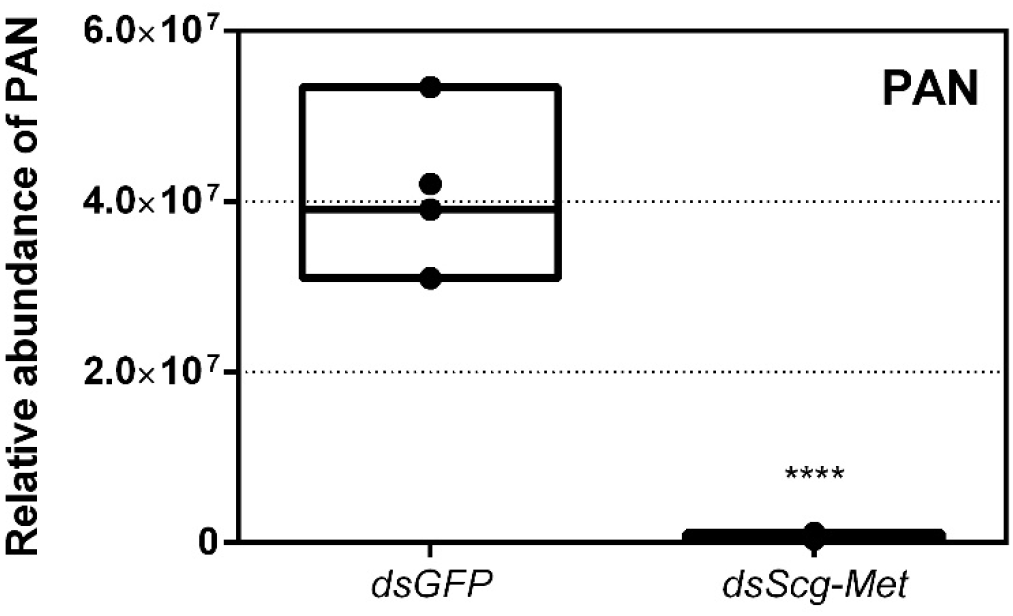
Relative abundance of PAN released by *dsGFP* and *dsScg-Met* injected adult male desert locusts 26 days after the adult molt. The emission of PAN was significantly reduced when *Scg-Met* was silenced. Data points are represented in a column graph as a floating bar (min to max value) with a line indicating the median. Significant differences (Student’s T-test) are indicated with asterisks (****, p < 0.0001). For each condition five animals were analyzed. PAN emission was measured by gas chromatography–mass spectrometry (GC–MS). **Abbreviations:** ds = double stranded, *Scg = Schistocerca gregaria*, GFP = Green Fluorescent Protein, Met = Methoprene-tolerant, PAN = phenylacetonitrile.

### Effect of *Scg-Met* knockdown on development of male reproductive organs

We also examined the effect of *Scg-Met* silencing on the weights of the testes and AG in adult male locusts. Thirty-four days after the first injections, the total body weight, as well as the weights of the testes and AG of both the *dsGFP* and the *dsScg-Met* treated animals were determined for the calculation of the GSI and AGSI, respectively (Fig. 5 A-C). At this point, none of the *dsScg-Met* injected males had initiated copulation with their female conspecifics, while on the other hand all control animals were sexually active. In line with this observation, animals subjected to a *Scg-Met* knockdown had significantly lower GSI (p = 0.0059) and AGSI (p < 0.0001) indices, when compared to the control animals (Fig. 5 B-C). Lower GSI and AGSI indicate lower normalized weights of both testes and accessory glands in the *Scg-Met* knockdown animals than in the controls. The impaired male reproductive organ development is further evidenced by the significantly lower normalized protein content of the AG in *dsScg-Met* injected locusts, when compared to the *dsGFP* control animals (p = 0.0255)(Fig. 5 D).

**Figure 5.**
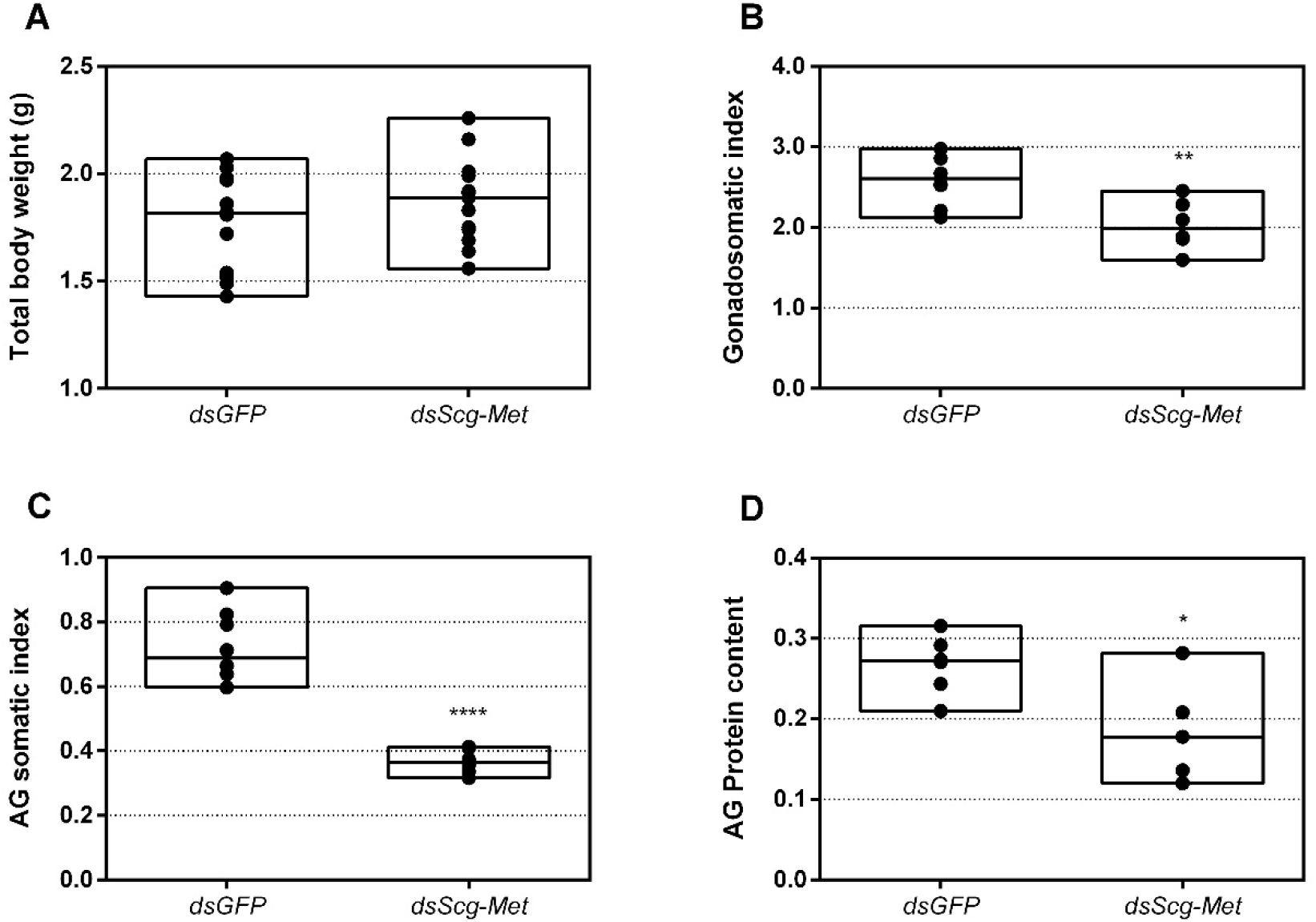
**(A)** Total body weight in grams of *dsGFP* and *dsScg-Met* injected adult male desert locusts 34 days after the adult molt. Data points are represented in a column graph as a floating bar (min to max value) with a line indicating the median. For each condition 12 animals were analyzed. No significant differences (Student’s T-test) in body weight were observed between both conditions (the statistical p-value obtained for total body weight was 0.2414). **(B)** Gonadosomatic index (GSI) of *dsGFP* and *dsScg-Met* injected adult male desert locusts 34 days after the adult molt. The GSI was significantly lower in *dsScg-Met* locusts compared to control animals. This result indicates the impaired development of the testes upon silencing of *Scg*-Met. Data points are represented in a column graph as a floating bar (min to max value) with a line indicating the median. For each condition 12 animals were analyzed. Significant differences (Student’s T-test) are indicated with asterisks (** p < 0.01) (the statistical p-value obtained for GSI was 0.0059). **(C)** Accessory gland somatic index (AGSI) of *dsGFP* and *dsScg-Met* injected adult male desert locusts 34 days after the adult molt. The AGSI was significantly lower in *dsScg-Met* injected locusts compared to *dsGFP* (control) animals. This result indicates the impaired development of the accessory glands (AG) upon silencing of *Scg*-Met. Data points are represented in a column graph as a floating bar (min to max value) with a line indicating the median. For each condition 12 animals were analyzed. Significant differences (Student’s T-test) are indicated with asterisks (**** p < 0.0001). **(D)** Normalized protein content of the accessory glands of *dsGFP* and *dsScg-Met* injected adult male desert locusts 34 days after the adult molt. The protein content of the AG was determined using the bicinchoninic acid assay (BCA) method and normalized to the respective AG weight. The relative protein content in the AG of *dsScg-Met* injected animals was significantly reduced compared to that of *dsGFP* injected (control) locusts. Data points are represented in a column graph as a floating bar (min to max value) with a line indicating the median. For each condition 12 animals were analyzed. Significant differences (Student’s T-test) are indicated with asterisks (* p < 0.05) (the statistical p-value obtained for relative protein content in AG was 0.0255). **Abbreviations:** ds = double stranded, *Scg = Schistocerca gregaria*, GFP = Green Fluorescent Protein, Met = Methoprene-tolerant.

### Effect of *Scg-Met* knockdown on gene expression profiles in the fat body

In this study, we also determined the levels of three developmental peptide hormone encoding transcripts upon *dsGFP* or *dsScg-Met* injection. Thirty-four days after the adult molt, we observed significantly elevated *Scg-NP3* (p = 0.0059) and *Scg-NP4* (p = 0.0007) and downregulated *Scg-IRP* (p = 0.0021) transcript levels in the fat body of *dsScg-Met* treated animals, when compared to the *dsGFP* injected control animals (Fig. 1 C-D-E). Interestingly, upon dissection, we also observed that the fat body in the *dsScg-Met* treated animals was more pronounced than in the *dsGFP* controls (Suppl. Fig. S2).

### Silencing *Scg-Tai* also results in an identical phenotype

Earlier research in *Aedes aegypti, T. castaneum* and *L. migratoria* described the interaction of JH-bound Met with Tai to become a functional JH receptor complex that mediates downstream JH signaling via Kr-h1 (Li et al., 2011; Z. Wang et al., 2017; Zhang et al., 2011). Therefore, we also decided to perform an RNAi mediated knockdown of *Scg-Tai* in a similar way as described for the *Scg-Met* knockdown experiment. Animals were treated in a similar way as during the *Scg-Met* knockdown experiment and were also sacrificed at day 34 after molting to the adult stage. Similarly as observed upon silencing *Scg-Met*, the *dsScg-Tai* injected males lacked yellow pigmentation in their cuticle and, in line with this, their *Scg-YP* transcript level was significantly reduced compared to *dsGFP* injected control animals (p = 0.037) (Fig. 6 A). In addition, they also showed a drastically reduced PAN pheromone emission (p = 0.0018)(Fig. 6 B). Furthermore, the *dsScg-Tai* injected animals had a significantly reduced GSI (p = 0.0001)(Fig. 6 C). Accordingly, none of the *Scg-Tai* knockdown males demonstrated copulation behavior, as opposed to all *dsGFP* injected control males. Moreover, similarly as described above for the *dsScg-Met* condition, we observed a significant increase in the normalized surface areas of the CA after *dsScg-Tai* treatment in comparison with the controls (p < 0.0001)(Fig. 6 D). On average, the normalized surface area of the CA in control animals was 26.83 µm^2^/mg body weight, while the normalized surface area of the CA in *dsScg-Met* treated animals was 80.3 µm^2^/mg body weight. Additionally, the fat body of the *dsScg-Tai* injected males was more prominent than in locusts of the control condition and had a similar appearance as observed in the *dsScg-Met* injected condition (Suppl. Fig. S2).

**Figure 6.**
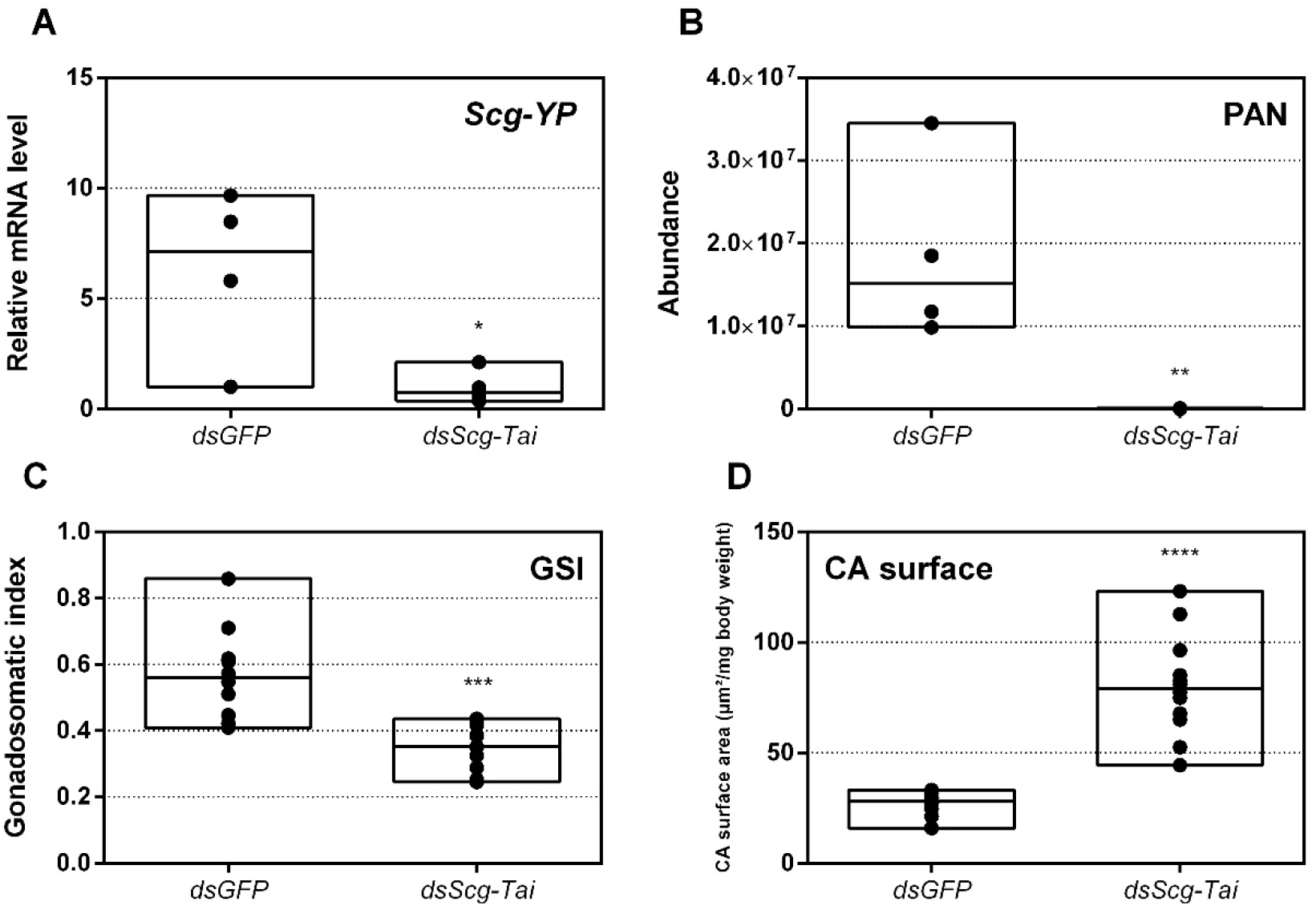
**(A)** Relative mRNA levels of *Scg-YP* measured in the epidermis of *dsGFP* and *dsScg-Tai* injected adult male desert locusts 34 days after the adult molt. *Scg-YP* transcript levels were significantly reduced after *Scg-Tai* silencing. Transcript levels in the epidermis were normalized against two reference genes, *Scg-Act* and *Scg-Ef1a*. Data points are represented in a column graph as a floating bar (min to max value) with a line indicating the median. For every condition five biological replicates, each consisting of epidermis samples from four individual locusts, were analyzed. Significant differences (Student’s T-test) are indicated with asterisks (* p < 0.05) (the statistical p-value obtained for *Scg-YP* was 0.0371). **(B)** Relative abundance of PAN released by *dsGFP* (control) and *dsScg-Tai* injected adult male desert locusts 26 days after the adult molt. The emission of PAN was significantly reduced when *Scg-Tai* was silenced. Data points are represented in a column graph as a floating bar (min to max value) with a line indicating the median. PAN emission was measured by gas chromatography–mass spectrometry (GC–MS). For every condition five animals were analyzed. Significant differences (Student’s T-test) are indicated with asterisks (** p < 0.01) (the statistical p-value obtained for PAN emission was 0.0018). **(C)** Gonadosomatic index (GSI) of control animals compared to *dsScg-Tai* treated animals. The GSI is significantly lower in *dsScg-Tai* treated animals compared to the control animals. This result indicates the impaired development of the testes upon silencing of *Scg-Tai*. Data points are represented in a column graph as a floating bar (min to max value) with a line indicating the median. For each condition 12 animals were analyzed. Significant differences (Student’s T-test) are indicated with asterisks (*** p < 0.001) (the statistical p-value obtained for GSI was 0.0001). **(D)** Normalized surface area of the CA in µm /mg body weight of *dsGFP* and *dsScg-Tai* treated animals. These data demonstrate that the CA significantly increased after *dsScg-Tai* treatment. On average, the normalized surface area of the CA in control animals was 26.83 µm^2^/mg body weight, while the normalized surface area of the CA in *dsScg-Met* treated animals was 80.3 µm^2^/mg body weight. Data points are represented in a column graph as a floating bar (min to max value) with a line indicating the median. For each condition 13 animals were analyzed. Significant differences (Student’s T-test) are indicated with asterisks (**** p < 0.0001). **Abbreviations:** ds = double stranded, *Scg = Schistocerca gregaria*, GFP = Green Fluorescent Protein, Tai = Taiman, YP = yellow protein, Act = actin, Ef1a = elongation factor 1-alpha.

## Discussion

The body of research focusing on the role of the JH receptor components, Met and Tai, in sexual maturation of male insects is out of balance with the vast amount of information available for female insects. In the females of several insect species, a stimulatory role of the JH signaling system on vitellogenin production by the fat body and on oocyte development in the ovary is evidenced (Badisco et al., 2013). Also in adult *S. gregaria* females, we recently described that the JH receptor Met is necessary for ovarian maturation, vitellogenesis and associated ecdysteroid biosynthesis, making it a crucial gene in female locust reproductive physiology (Gijbels et al., 2019). In the current study, we report on the effects of an RNAi-mediated knockdown of *Scg-Met* and *Scg-Tai* on the reproductive development of adult male desert locusts. Our observations from these RNAi experiments indicate that in male *S. gregaria* both JH receptor components are essential for sexual maturation.

*Scg-Met* mRNA levels were significantly reduced by injecting the locusts with *dsScg-Met* indicating that the RNAi procedure was effective (Fig. 1 A). In addition, the levels of *Scg-Krh1* and *Scg-E93*, situated downstream of Met in the MEKRE93 pathway in juvenile insects, were investigated as well (Belles, 2020; Belles and Santos, 2014; Lozano and Belles, 2011; Ureña et al., 2014). The significant reduction in *Scg-Krh1* demonstrates that JH signaling was impaired upon silencing *Scg*-Met (Fig. 2 A). E93 is an ecdysone inducible transcription factor, of which the expression is inhibited by Kr-h1, and *vice versa* (Belles and Santos, 2014; Ureña et al., 2014). In line with this, an increase in the relative mRNA levels of *Scg*-E93 was observed in the CA (Fig. 2 A). Therefore, the contrasting expression profiles of *Scg-Krh1* and *Scg-E93* are very well in agreement with the sustained *Scg-Met* knockdown that was obtained by the regular *dsScg-Met* injections, suggesting that a mutually inhibiting cross-talk between *Scg*-Kr-h1 and *Scg*-E93 also persists in adult male locusts.

Silencing of *Scg-Met* resulted in very strong phenotypic effects, indicating a pivotal role of the JH signaling pathway in the life history of male desert locusts:

1. An obvious visual effect of the *Scg-Met* knockdown was the impaired cuticular pigmentation of the adult males (Fig. 3). *S. gregaria* adults are weakly pigmented after their final molt. Hardening of the cuticle upon molting, results in steady background coloration. In adult gregarious males, a remarkable change to a bright yellow cuticular coloration occurs when they become sexually mature (Uvarov, 1966). In our colony, this color change normally manifests itself at days 9-12 after the final molt. Research has highlighted the role of yellow protein (YP) in this remarkable phenotypic transformation. YP has a high binding capacity for carotenoids originating from the diet and will be allocated towards the epidermis and cuticle when gregarious males become sexually mature. High levels of JH during adult male development will induce increasing YP levels, eventually resulting in the deposition of the yellow pigment in the cuticle (Wybrandt and Andersen, 2001). The strongly reduced *Scg-YP* expression in *Scg-Met* silenced males is fully in line with the observed absence of a bright yellow coloration in the cuticle, in stark contrast with the *dsGFP* injected (control) males (Fig. 3).
2. In comparison to *dsGFP* injected (control) males, the *Scg-Met* knockdown locusts produced much lower amounts of the pheromone PAN (Fig. 4). Gregarious desert locusts live in very dense populations. Besides visual cues, pheromone communication plays a crucial role in mating behavior. One of the best-known pheromones produced by gregarious male desert locusts is phenylacetonitrile (PAN). This pheromone is abundantly released by the legs and wings of gregarious male locusts and plays a crucial role in identifying an appropriate sexual partner by preventing male-male interactions, as well as in the post-copulatory guarding of the inseminated female mate (Amwayi et al., 2012). Silencing of *Scg-Met* very strongly affected PAN emission, which illustrates that JH signaling plays a crucial role in the emission of this male-to-male anti-aphrodisiac pheromone.
3. The development of the male reproductive organs, testes and accessory glands, was significantly inhibited by silencing *Scg-Met* during the adult stage (Fig. 5). Interestingly, a recent study demonstrated that treating male *S. gregaria* nymphs (final nymphal stage) with a JH mimic resulted in underdeveloped accessory glands and seminal vesicles, and impaired molting to the adult stage (Hiroyoshi et al., 2019). This suggests a different function of JH in the development of reproductive organs in the desert locust during distinct life stages, again highlighting the complex role of JH throughout the life history of insects. The role of JH in protein accumulation in AG has been described in several insect species (Chen, 1984). A more recent study on *D. melanogaster Met*^27^ mutants first demonstrated the role of Met in the development of the male AG. *Met*^27^ mutants showed reduced AG sizes. This could be ascribed to the overall reduced protein synthesis resulting from the impaired JH signaling (Wilson et al., 2003). Similar effects were observed after knocking down Met or Tai in the linden bug, *Pyrrhocoris apterus* (Hejnikova et al., 2016). In the red flour beetle, *T. castaneum*, a JHAMT knockdown reduced the AG size and production of seminal proteins, significantly affecting male fitness (Parthasarathy et al., 2009). Also, in locust species, JH has previously been shown to be essential for the growth of the AG and associated protein content in this tissue (Avruch and Tobe, 1978; Braun and Wyatt, 1995; Ismail and Gillott, 1996). Our results show that JH receptor silencing in *S. gregaria* produced effects opposite to JH itself, clearly illustrating the adverse consequences of a disabled transduction of the JH signal on the development of reproductive organs in adult male locusts.
4. Interestingly, both the size of the CA and the expression of *Scg-JHAMT*, which codes for a rate-limiting JH biosynthetic enzyme, were significantly increased upon sustained silencing of *Scg-Met* (Fig. 2). These observations suggest the existence of homeostatic regulatory mechanisms for preserving a balance between JH synthesis and JH sensitivity within the CA. A similar stimulation of JH production following Met knockdown was previously also described in the linden bug, *P. apterus* (Hejnikova et al., 2016).
5. Finally, silencing of either component of the JH receptor complex induced very similar phenotypic effects in the desert locust (Fig. 6). Recent studies revealed that Met forms a complex with Tai to become a functionally active JH-dependent receptor. Once JH is bound to Met, the JH-Met complex will subsequently interact with a Tai transcription factor domain to form an active JH signal transducer (Charles et al., 2011). Our findings are in line with this mechanistic model for JH signaling and indicate that reproductive organ growth, yellow coloration and PAN pheromone emission in gregarious males are highly dependent on expression of both *Scg-Met* and *Scg-Tai*.

In this study, we have also analyzed the transcript profiles of *Scg-Met*, its dimerization partner *Scg-Tai* and the downstream factor *Scg-Krh1* during male adult development in four different tissues and on three different time points after the final molt (Suppl. Fig. S1). The abundance of the transcripts coding for these JH pathway components significantly increased in the fat body during male reproductive maturation, showing lowest levels when males were sexually immature (day 3), intermediate levels when they were maturing (day 7) and highest when they had reached sexual maturity (day 15) (Suppl. Fig. S1). On day 15, males in our gregarious desert locust colony have fully developed gonads, have a bright yellow colored cuticle and are sexually active as manifested by their mating behavior. That the levels of all three transcripts significantly increased from day 3, over day 7 to day 15 in male fat body indicates that JH directly interacts with male fat body. This increased expression of receptor components in the fat body seems to correlate with the significant rise of mRNA levels for the JH biosynthetic enzymes *Scg*-JHAMT and *Scg*-CYP15A1 in the CA, both crucial for JH biosynthesis. This is also in line with the observed increase in *Scg-Krh1* expression, which is known to be activated downstream of the interaction of JH with Met and of JH-bound Met with Tai. These data may point at a prominent role of the fat body in generating JH-dependent responses that alter the locust’s metabolism in order to regulate the physiological development of male reproductive organs. Interestingly, in CA/CC complexes the transcript levels of *Scg-Met, Scg-Tai* and *Scg-Krh1* were significantly lower on days 7 and 15 when compared to day 3. Together with findings represented in Fig. 2 and Fig. 6, which show the significantly increased *Scg-JHAMT* expression, as well as larger CA size, upon *Scg-Met* or *Scg-Tai* knockdown, this observation strongly suggests the existence of homeostatic control mechanisms for inversely regulating JH biosynthesis and JH signaling activity within the CA.

Considering the important role of the fat body in metabolic, developmental and reproductive regulation, we have investigated the levels of *Scg-IRP, Scg-NP3* and *Scg-NP4*, three important peptide hormone encoding transcripts that are known to be expressed in the fat body and play a role in female reproductive physiology (Badisco et al., 2011; Claeys et al., 2003). When compared to *dsGFP* injected control males, the *dsScg-Met* injected ones had significantly reduced *Scg-IRP* levels in the fat body (Fig. 1 C). In many metazoans, the insulin/IGF signaling pathway (ISP) is an important sensor of the nutritional and metabolic status, as well as a regulator of anabolic processes that result in growth and/or reproduction (Badisco et al., 2013; Nässel and Vanden Broeck, 2016). In *S. gregaria*, only one insulin-related peptide (IRP) (*Scg*-IRP) with main expression in brain and fat body was identified (Badisco et al., 2008). Previous studies revealed that *Scg*-IRP expression is regulated during the reproductive cycle and there is evidence for a stimulatory role of *Scg*-IRP on vitellogenesis and ovarian development in adult female desert locusts (Badisco et al., 2011). In several other insect species, complex functional interactions have been observed between the ISP and JH signaling (Badisco et al., 2013; Gijbels et al., 2019; Roy et al., 2018; Van Wielendaele et al., 2012). In the fruit fly, *D. melanogaster*, the expression of the insulin receptor (IR) was detected in the CA, indicating the ability of the CA to respond to changing insulin-like peptide levels. Moreover, *D. melanogaster IR* mutants showed significantly reduced JH levels (Tatar et al., 2001). In the mosquito *Culex pipiens*, the administration of JH rescued the defects observed upon silencing of the ISP (Sim and Denlinger, 2009). A microarray analysis to explore gene expression in response to the administration of a JH analog in the silkworm, *Bombyx mori*, demonstrated a stimulatory effect on key regulatory genes involved in the ISP (Cheng et al., 2014). In the red flour beetle, *T. castaneum*, JH signaling was shown to stimulate the expression of two insulin-like peptide encoding genes (Xu et al., 2013). Moreover, in this species, silencing of the JH receptor Met resulted in decreased insulin-like peptide transcript levels (Sheng et al., 2011). Therefore, our data of the current study extend these widely observed links between JH signaling and ISP towards reproductive development of adult male desert locusts. In this process, an important role seems to be played by the fat body, where the expression of *Scg-Met, Scg-Tai* and *Scg-Krh1* significantly increased during male sexual maturation (Suppl. Fig. S1), and where in *dsScg-Met* injected males significantly reduced transcript levels were observed for both *Scg-Met* and *Scg-IRP* (Fig. 1). In addition, the fat body appeared to be more pronounced in both *Scg-Met* and *Scg-Tai* knockdown males than in the *dsGFP* injected (control) ones (Suppl. Fig. S2). Interestingly, fat body hypertrophy is generally observed in insects with reduced JH and ISP signaling, and in several species this is a naturally occurring phenomenon when the animal is going into a reproductive diapause (Badisco et al., 2013). This happens in situations when the environmental conditions are adverse, and energy and nutrients are rather saved for the individual insect’s survival than invested in gonad development and reproduction. Moreover, it has been shown that a reproductive diapause can be artificially induced in the mosquito *C. pipiens* by knocking down the insulin receptor via RNAi (Sim and Denlinger, 2009).

While the levels of *Scg-Met* and *Scg-IRP* mRNAs in the fat body of *dsScg-Met* injected males were reduced, we observed significantly elevated expression of *Scg-NP3* and *Scg-NP4*. In locusts, NPs were originally characterized based on their antigonadotropic effects in female desert locusts (Badisco et al., 2011). The effects generated by NPs are opposite to these of JH and IRP, although the exact *in vivo* relationship with these pathways remains elusive. Four closely related members of the neuroparsin family have previously been cloned from *S. gregaria* (Badisco et al., 2011; Claeys et al., 2003; Janssen et al., 2001), but mRNA levels of only two (*Scg-NP3* and *Scg-NP4*) are predominantly localized in the fat body of gregarious desert locusts. In locusts, the levels of these neuroparsin transcripts show temporal changes during the hormonally controlled molting and reproductive cycles, while locust phase dependent differences in NP expression have also been reported (Claeys et al., 2006, 2003). Interestingly, in line with the observed sequence similarity of NPs with a highly conserved region in vertebrate IGBP, *Scg*-NP4 was shown capable of interacting *in vitro* with *Scg*-IRP (Badisco et al., 2008). When knocking down all four neuroparsins in female locusts, a clear increase was observed in vitellogenin transcript levels in the fat body and in basal oocyte size in the ovarian follicles (Badisco et al., 2011). In the present study, we observed a remarkably opposite effect of *Scg-Met* silencing on *Scg-NP3/4* expression in comparison to *Scg-IRP*. Increased NP and decreased IRP levels would be expected to strengthen the inhibitory effects on reproductive development imposed by silencing *Scg*-Met, the receptor responsible for JH signaling. Therefore, the changes in *Scg-IRP, Scg-NP3* and *Scg-NP4* expression observed after *Scg-Met* knockdown may have contributed to the arrested development of the male reproductive system, the inability to acquire mature gregarious male traits and the observed sexually inactive phenotype.

In conclusion, this study has demonstrated that the proper development of sexual traits in adult gregarious locust males can be inhibited in an early stage by injecting the locusts as of adult molting with either *dsScg-Met* or *dsScg-Tai*. The treatment resulted in individuals that showed no mating behavior, did not produce the associated pheromone and did not obtain the characteristic bright yellow color associated with sexually mature male locusts. Our results therefore indicate a “master switch” role of these JH receptor components in sexual development of male *S. gregaria*. Our results clearly demonstrate the potential of manipulating locust male reproductive maturation to control locust population growth. Specifically targeting the JH signaling system of locusts could therefore prove to be a very efficient and successful way by which swarming desert locust plagues can be prevented in a more biorational manner.

## Acknowledgements

The authors would like to thank Evert Bruyninckx for technical assistance and Evelien Herinckx for taking care of the locust rearing facility. This work was supported by grants from the Special Research Fund of KU Leuven (C14/15/050; C14/19/069), the European Union’s Horizon 2020 Research and Innovation programme (No. 634361), and the Research Foundation of Flanders (FWO) (G0F2417N; G090919N) to JVB. MH was supported by a PhD fellowship from the Agency for Innovation by Science and Technology (IWT); MG and JVL obtained a PhD fellowship from the Fund for Scientific Research (FWO).

## Author contributions

Experiments with locusts were designed by MH and EM and performed by MH, MG, JVL and EM. The study was supervised by EM and JVB. Measurements of the locust volatiles were performed by ED under supervision of BN. The initial version of the manuscript was written by MH, EM and JVB, and further edited by all authors. JVB is the principle investigator responsible for funding acquisition and project administration.

## Declaration

The authors declare no conflict of interest, financial or otherwise, that might potentially bias this work.

---

**Supplementary Figure S1.**
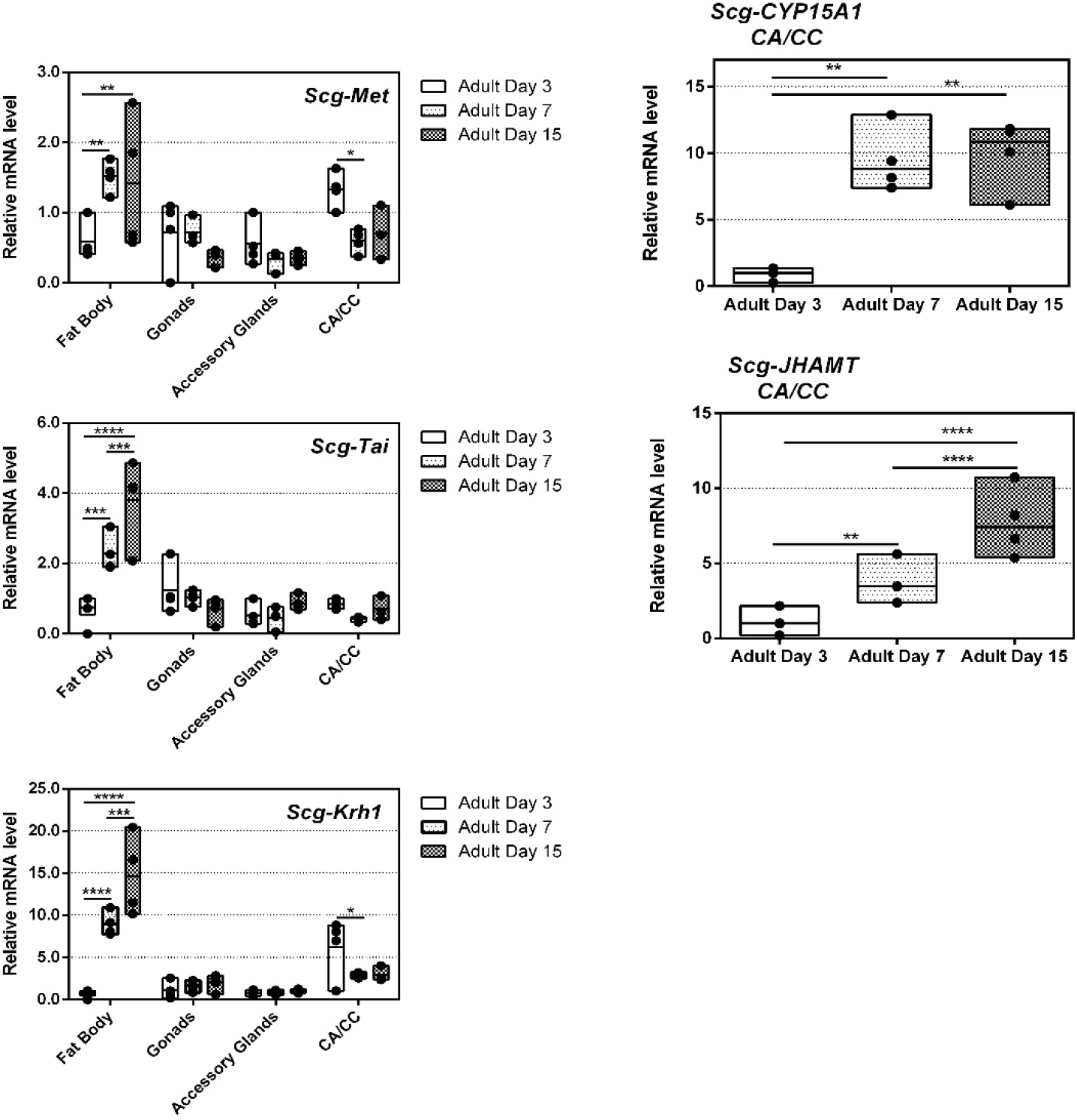
Relative mRNA levels of *Scg-Met, Scg-Tai* and *Scg-Krh1* measured in the fat body, gonads, accessory glands and CA/CC of male desert locusts at day 3, day 7 and day 15 following the adult molt. Relative mRNA levels of *Scg-JHAMT* and *Scg-Cyp15* were only measured in the CA/CC complexes, since the CA are the primary site of synthesis and activity for both enzymes. Transcript levels of *Scg-Met, Scg-Tai* and *Scg-Krh1* significantly increased from day 3, over day 7, to day 15 in the fat body of adult male desert locusts However, no significant changes in the levels of these transcripts were detected in either the testes or the accessory glands. In the CA/CC complex of male desert locusts, the relative transcript levels of *Scg-JHAMT* and *Scg-CYP15A1* significantly increased from day 3 to days 7 and 15 of the adult stage. In the CA/CC, the transcript levels of *Scg-Met* and *Scg-Krh1* were significantly lower on days 7 and 15, when compared to day 3. For every condition four biological replicates, each consisting of the pooled tissues from five individual locusts, were analyzed by q-RT-PCR. Transcript levels were normalized against two reference genes, *Scg-Act* and *Scg-Ef1a*. Data points are represented in a column graph as a floating bar (min to max value) with a line indicating the median. Significant differences (ANOVA) are indicated with asterisks (* p < 0.05; ** p < 0.01; *** p < 0.001; **** p < 0.0001). The following statistical p-values were obtained: for *Scg-Met* in the fat body: day 3 vs day 7 (p = 0.0036), day 3 vs day 15 (p = 0.0097), day 7 vs 15 (p = 0.9249); in the gonads: day 3 vs day 7 (p = 0.9997), day 3 vs day 15 (p = 0.4120), day 7 vs 15 (p = 0.3999); in the accessory glands: day 3 vs day 7 (p = 0.7108), day 3 vs day 15 (p = 0.7446), day 7 vs 15 (p = 0.9983); in the CA/CC: day 3 vs day 7 (p = 0.0249), day 3 vs day 15 (p = 0.0921), day 7 vs 15 (p = 0.9227). For Scg-Tai in the fat body: day 3 vs day 7 (p = 0.0009), day 3 vs day 15 (p = < 0.0001), day 7 vs 15 (p = 0.0003); in the gonads: day 3 vs day 7 (p = 0.8207), day 3 vs day 15 (p = 0.2850), day 7 vs 15 (p = 0.6191); in the accessory glands: day 3 vs day 7 (p = 0.9759), day 3 vs day 15 (p = 0.6211), day 7 vs 15 (p = 0.4922); in the CA/CC: day 3 vs day 7 (p = 0.4767), day 3 vs day 15 (p = 0.9253), day 7 vs 15 (p = 0.7588). For *Scg-Krh1* in the fat body: day 3 vs day 7 (p = < 0.0001), day 3 vs day 15 (p = < 0.0001), day 7 vs 15 (p = 0.0004); in the gonads: day 3 vs day 7 (p = 0.9098), day 3 vs day 15 (p = 0.7653), day 7 vs 15 (p = 0.9563); in the accessory glands: day 3 vs day 7 (p = 0.9967), day 3 vs day 15 (p = 0.9891), day 7 vs 15 (p = 0.9978); in the CA/CC: day 3 vs day 7 (p = 0.0473), day 3 vs day 15 (p = 0.0769), day 7 vs 15 (p = 0.9991), For Scg-CYP15A1 in the CA/CC: day 3 vs day 7 (p = 0.0024), day 3 vs day 15 (p = 0.0018), day 7 vs 15 (p = 0.9560). For *Scg-JHAMT* in the CA/CC: day 3 vs day 7 (p = 0.0011), day 3 vs day 15 (p < 0.0001), day 7 vs 15 (p < 0.0001).

**Supplementary Figure S2.**
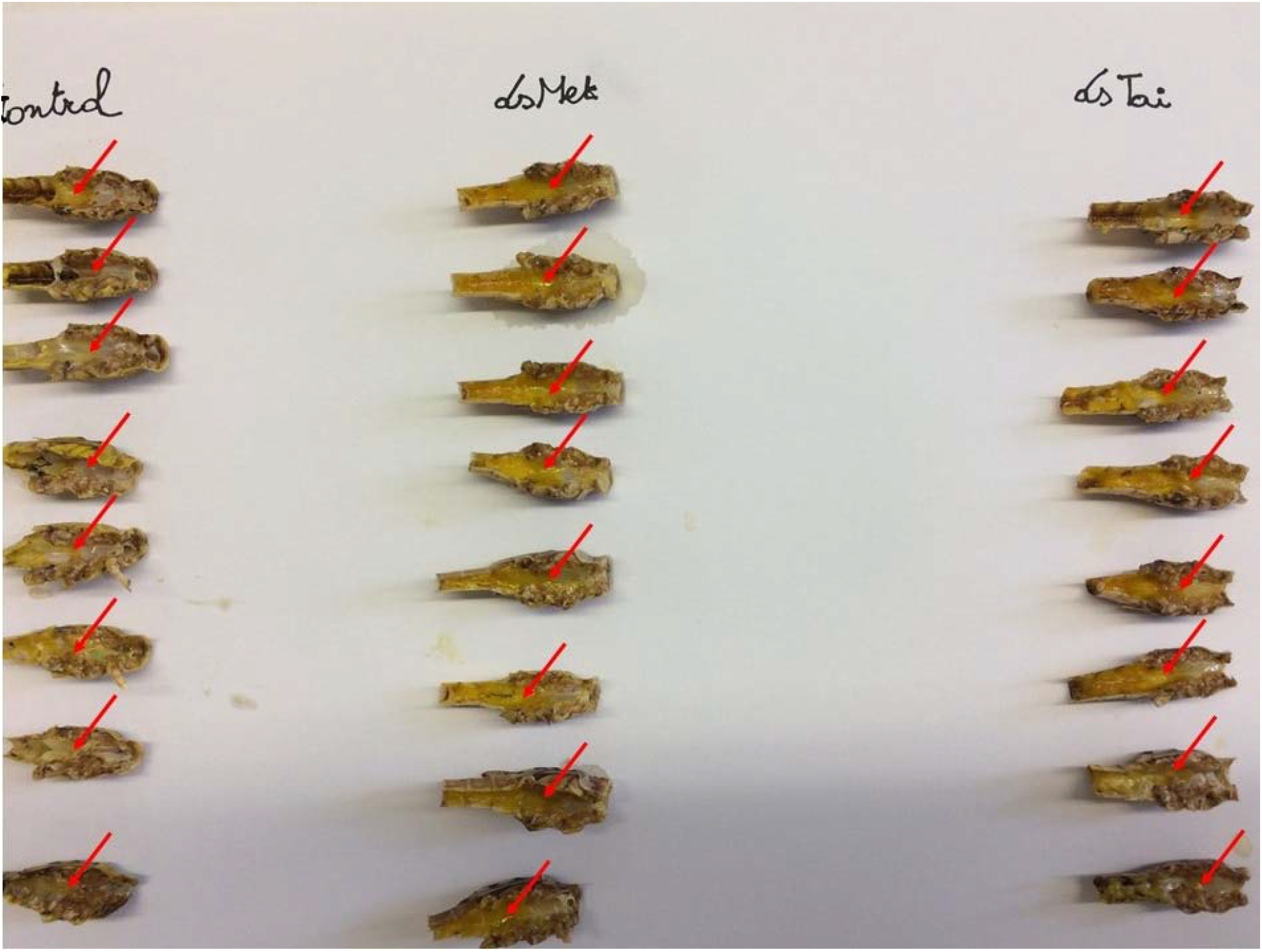
Dissected carcasses of *dsGFP* (control), *dsScg-Met* (dsMet) and *dsScg-Tai* (dsTai) injected adult male *Schistocerca gregaria*. These pictures clearly illustrate that the fat body in the *dsScg-Met* and *dsScg-Tai* injected males was more pronounced than in animals of the *dsGFP* injected control condition. The fat body is visible as the yellow-beige tissue in the carcasses of the dissected locusts, as indicated in the figure with red arrows.

